# Exploring the Free Energy Landscape and Thermodynamics of Protein-Protein Association: HIV-1 Integrase Multimerization Induced by an Allosteric Inhibitor

**DOI:** 10.1101/2020.06.29.177592

**Authors:** C. Tse, L. Wickstrom, M. Kvaratskhelia, E. Gallicchio, R. Levy, N. Deng

## Abstract

We report the free energy landscape and thermodynamics of the protein-protein association responsible for the drug-induced multimerization of HIV-1 integrase (IN). Allosteric HIV-1 integrase inhibitors (ALLINIs) promote aberrant IN multimerization by bridging IN-IN intermolecular interactions. However, the thermodynamic driving forces and kinetics of the multimerization remain largely unknown. Here we explore the early steps in the IN multimerization by using umbrella sampling and unbiased molecular dynamics simulations in explicit solvent. In direct simulations, the two initially separated dimers spontaneously associate to form near-native complexes that resemble the crystal structure of the aberrant tetramer. Most strikingly, the effective interaction of the protein-protein association is very short-ranged: the two dimers associate rapidly within tens of nanoseconds when their binding surfaces are separated by d ≤ 4.3 Å (less than two water diameters). Beyond this distance, the oligomerization kinetics appears to be diffusion controlled with a much longer association time. The free energy profile also captured the crucial role of ALLINI in promoting multimerization, and explained why several CTD mutations are remarkably resistant to the drug-induced multimerization. The results also show that at small separation the protein-protein binding process contains two consecutive phases with distinct thermodynamic signatures. First, inter-protein water molecules are expelled to the bulk resulting in a small increase in entropy, as the solvent entropy gain from the water release is nearly cancelled by the loss of side chain entropies as the two proteins approach each other. At shorter distances, the two dry binding surfaces adapt to each other to optimize their interaction energy at the expense of further protein configurational entropy loss. While the binding interfaces feature clusters of hydrophobic residues, overall, the protein-protein association in this system is driven by enthalpy and opposed by entropy.

**Statement of Significance:** Elucidating the energetics and thermodynamic aspects of protein-protein association is important for understanding this fundamental biophysical process. This study provided a more complete physical picture of the protein-protein association responsible for the drug-induced HIV-1 integrase multimerization. The results captured the critical role of the inhibitor, and accounted for the effects of mutations on the protein association. Remarkably, the effective range of the protein-protein attractive funnel is found to be very short, at less than two layers of water, despite the fact that the two binding partners carry opposite net charges. Lastly, entropy/enthalpy decomposition shows that the solvent release from the inter-protein region into the bulk is more than offset by the loss of the solute configurational entropy due to complexation.

## Introduction

Noncovalent and specific protein-protein association plays a crucial role in a wide range of fundamentally important biological processes, such as signal transduction and antibody-antigen recognition^1^. In recent years, protein-protein interactions are increasingly targeted for therapeutics development^2–3^. In this work, we characterize at an atomic level of detail the free energy landscape and the thermodynamic driving forces for the protein-protein association underlying the drug-induced HIV-1 integrase multimerization^4–7^. HIV-1 integrase (IN) is an important therapeutic target for the development of antiviral therapy^2^, because of the vital roles IN plays in the life cycle of HIV. Allosteric HIV-1 integrase inhibitors (ALLINIs) bind at the dimer interface of the IN catalytic core domain (CCD) and inhibit HIV-1 by promoting aberrant IN multimers, which impairs both the catalytic activity of IN and virus core maturation.^8–10^ The mechanism by which ALLINIs promote aberrant multimerization has previously been revealed by our protein-protein docking simulations which suggested that the multimerization is mediated by ALLINI which bridges IN-IN interactions between the catalytic core domains (CCD) of one IN dimer and the C-terminal domain (CTD) of another dimer. The central prediction from the modeling study of the IN aberrant multimerization, i.e. the CCD and CTD intersubunit interaction bridged by ALLINI^5^, was confirmed by the experimental crystal structure later obtained containing full-length IN and ALLINI^11^.

Despite the progress in understanding the ALLINI-induced aberrant multimerization, several important aspects of the molecular mechanism of the aberrant multimerization need to be better understood in order to inform rational inhibitor design. What is the predominant thermodynamic driving force for the intermolecular association of CTD and CCD bridged by ALLINI? How does the association free energy landscape of the IN multimerization look like? At the molecular level, why do specific amino acid substitutions in the CTD such as Y226D and K266A block the ALLINI-induced multimerization? What are the roles of the water molecules in the intervening space between the CCD and CTD in the association? To date, only limited, indirect experimental data regarding the drug-induced IN multimers are available. The only experimental structure of an aberrant multimer was determined at a relatively low resolution of 4.4 Å^11^. While our modeling correctly predicted the central mechanism of action of the drug-induced multimerization, the use of proteinprotein docking precludes the characterization of the association mechanism directly influenced by solvent.

To address the mechanistic questions regarding the IN multimerization at an atomic level of detail, here we employ molecular dynamics (MD) simulations in explicit solvent to study the CCD and CTD association mediated by ALLINI. In recent years, because of the advances in methodology, algorithm and computer hardware, MD simulation and associated techniques have increasingly been successfully used to study the energetics and kinetics of protein-protein binding^12–15^. Here, we first directly simulate the formation of a tetramer starting from two dimers initiated from dissociated states with different separations between the CTD and CCD. The results show that the experimental multimer structure can be obtained using unbiased physical models in explicit aqueous solutions. We then quantitatively explore the free energy landscape of the protein-protein association using umbrella sampling with varying conditions to investigate the role of ALLNI and the different amino acid mutations that influence the multimerization. Entropy/enthalpy decomposition of the free energy reveals a detailed thermodynamic picture of protein-protein association that is dominated by favorable overall enthalpy, despite the presence of hydrophobic interactions traditionally believed to be favored by entropy. The study provides a wealth of atomistic information concerning the biophysics of drug-induced IN aberrant multimerization that may inform the design of new allosteric IN inhibitors.

## Materials and Methods

The GROMACS program 2016-3^16–17^ was used to performed the MD simulations in this work. The starting structure of the IN dimers (Fig. 1) was taken from the structure of the aberrant tetramer containing the ALLINI BI224436 described in the previous report^4^. The protein was modeled by the AMBER parm99ILDN force field^18^, while the ligand by the Amber GAFF^19^ and the AM1-bcc charge model^20^. Truncated octahedral boxes containing TIP3P water^21^ were used to solvate the solute, with the distance between any solute atom and the nearest wall of the box is ≥ 10 Å. Sodium and chloride ions are added to the solvent box to neutralize the net charges on the solute and to mimic the experimental ionic concentration of 0.1 M. The electrostatic interactions were treated using the particle-mesh Ewald (PME)^22^ method with a real space cutoff of 10 Å and a grid spacing of 1.0 Å. MD simulations were performed in the NPT ensemble with a time step of 2 fs.

**Figure 1.**
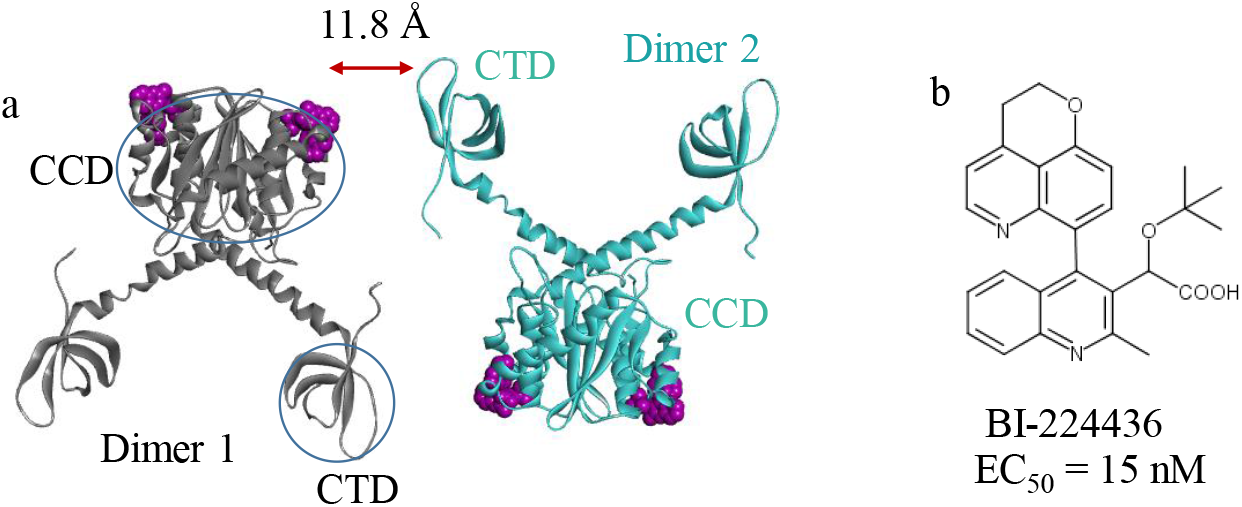
a. Two HIV-1 integrase dimers separated by interfacial gap d = 11.8 Å are solvated in a box of water (not shown). b. Chemical structure of the ALLINI BI224436.

The potentials of mean force were computed using umbrella sampling by applying harmonic biasing potential on the distance between the center of mass (COM) of the CTD (denoted as L1) and center of mass of the following CCD residues (denoted here as P1): 124-128, 131-132, 167-168 and BI224436 (Fig. 5). During the umbrella sampling simulation, weak angular harmonic restraints are also applied to maintain the relative orientation between the two proteins and to ensure that the pulling is done along the predefined axis ***r***_*P*1−*L*1_, and to accelerate the convergence of the PMF calculation. For this purpose, two polar angles (θ, ϕ) and three Euler angles (Θ, Φ, Ψ)^23–24^ are defined using the two centers of masses P1 and L1, plus two atoms on each of the binding proteins. The following two CCD atoms (P2, P3) and two CTD atoms (L2, L3) are chosen together with P1 and L1 to define five angles (*θ, ϕ*, Θ, Φ, Ψ) that determine the ligand orientation: P2: T124-CA; P3: W131-C; L2: D253-CA; L3: V234-C. The range of the pulling distance is 11.0 Å ≤ |***r***_*P*1−*L*1_| ≤ 23.9 Å. The equilibrium values of the five angles are: *θ*_0_ = 58.29°, *ϕ*_0_ = −70.92°, Θ_0_ = 130.1°, Φ_0_ = 77.27°, Ψ_0_ = −57.45°. A single force constant *k_r_* = 23.9 kcal mol^−1^ Å^−2^ is used for the distance restraints in all the umbrella sampling windows. The force constants used in the angular restraints are:*k_θ_* = *k_ϕ_* = *k*_Θ_ = *k*_Φ_ = *k*_Ψ_ = 0.0728 *kcal degree*^−2^*mol*^−1^.

For umbrella sampling used to compute the 1D PMF, 24 umbrella windows are used to cover the full range of the distance space. The spacing between adjacent umbrella windows is 0.5 - 1.0 Å, which we find provide good overlap between the sampled distances. In each umbrella window, a reasonable starting configuration is generated by first running a short 1.2 ns umbrella sampling simulation starting from the last snapshot of the short umbrella sampling simulation in the previous umbrella window. Next, 30 ns umbrella sampling simulations are run in parallel in each window starting from the initial configurations generated above. The last 20 ns trajectory was used to compute the PMF using the WHAM^25^ program implemented in the GORMACS 2016.3 package. For each protein-protein complex, four independent umbrella sampling simulations were run for estimating the statistical uncertainties in the calculated PMF. Thus, a total of ~2.88 μs simulation time is used to compute the PMF for a single protein-protein complex. The binding entropy is estimated using the temperature difference of the free energy changes at 275 K, 300 K and 320 K. The error bar in the Δ*S* is estimated using the formula for the standard error of the slope of linear regression, i.e. 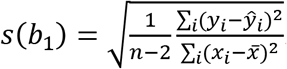. The error bar in the Δ*H* is estimated using error propagation from the statistical error in the *T*Δ*S* and Δ*G*.

## Results

### Direct simulation of the formation of aberrant tetramer from dimers

In a previous report, we have used protein-protein docking guided by indirect experimental information from hydrogen-deuterium exchange (HDX) data as restraints to derive the first structural model of the drug-induced HIV-1 integrase multimer^5^. The model accurately predicted that the intermolecular CCD-CTD interaction mediated by bound ALLINIs is the most important feature of the drug-induced IN multimerization. While the structural model predicted using protein-protein docking was confirmed by the subsequently reported crystal structure^11^, the docking study was carried out without the aqueous environment and therefore precluded the characterization of the thermodynamics and kinetics of the drug-induced IN aggregation. To explore the association mechanism of the IN multimerization process, here we perform multiple independent MD simulations in explicit solvent without any restraints for a total of 23 μs starting from two dissociated HIV-1 integrase dimers containing ALLINI BI224436^26–27^ with varying starting interfacial separations *d* (Fig. 1). The starting structure of two dimer is taken from the structure of the aberrant tetramer containing BI224436 reported previously^4^. The starting interface gap distance *d* is obtained by increasing the dimer-dimer separation from the structure of the aberrant tetramer complex by *d*. The setup and results of these simulations are summarized in Table 1. In simulation set-1, the initial separation distance between the surfaces of CTD and CCD of the two dimers is *d* ≈ 4.3 Å, while in the simulation set-2, they are initially separated by between 7.8 Å and 11.8 Å. All the simulations are repeated with different initial velocities and each lasts on average 500 ns. The structures sampled in the MD trajectories are compared with the crystal structure of the aberrant IN tetramer reported by Gupta *et al^11^*.

**Table 1.** Summary of the direct simulations of aberrant tetramer formation starting from dissociated IN dimers.

| Simulation Set | Starting Configuration | Successful association/total trajectories | MFPT<sup>a</sup> (ns) | Average trajectory length (ns) |
| --- | --- | --- | --- | --- |
| Set 1 | $d \approx 4.3$ Å | 8/10 | ~20 | 500 |
| Set 2 | $7.8 \text{ Å} \leq d \leq 11.8 \text{ Å}$ | 0/10 | > 500 | 500 |

a. Mean First Passage Time to the formation of near-native complex.

### The experimental aberrant tetramer structure is captured in unbiased MD simulations in explicit solvent starting from separated integrase dimers

We found that in the majority of the trajectories in Set 1, the two initially separated dimers (d ≈ 4.3 Å) spontaneously associate to form complexes that resemble the experimental structure of the native aberrant multimer: see Fig. 2, which compares a representative MD snapshot with the experimental structure. A snapshot is considered to have a near native multimer interface if the following criteria is met: when the central interface residues in the CCD (T124, A128, E170, H171 and T174) are superimposed onto the corresponding residues of the CCD in the crystal structure, the root mean square deviation (IRMSD) between the interface residues of the CTD (Y226, W235, K266 and I268) in the MD structure and those in the crystal structure is less than 3.5 Å. Importantly, the outcome of the simulation is highly sensitive to the starting separation of the two dimers: in contrast to the trajectories in Set 1, none of the trajectories in the Set 2 leads to successful association and the formation of aberrant tetramer. Note that the only difference between the two sets of simulations is that the two dimers in Set 2 have larger initial separation compared with those in Set 1 (Table 1).

**Figure 2.**
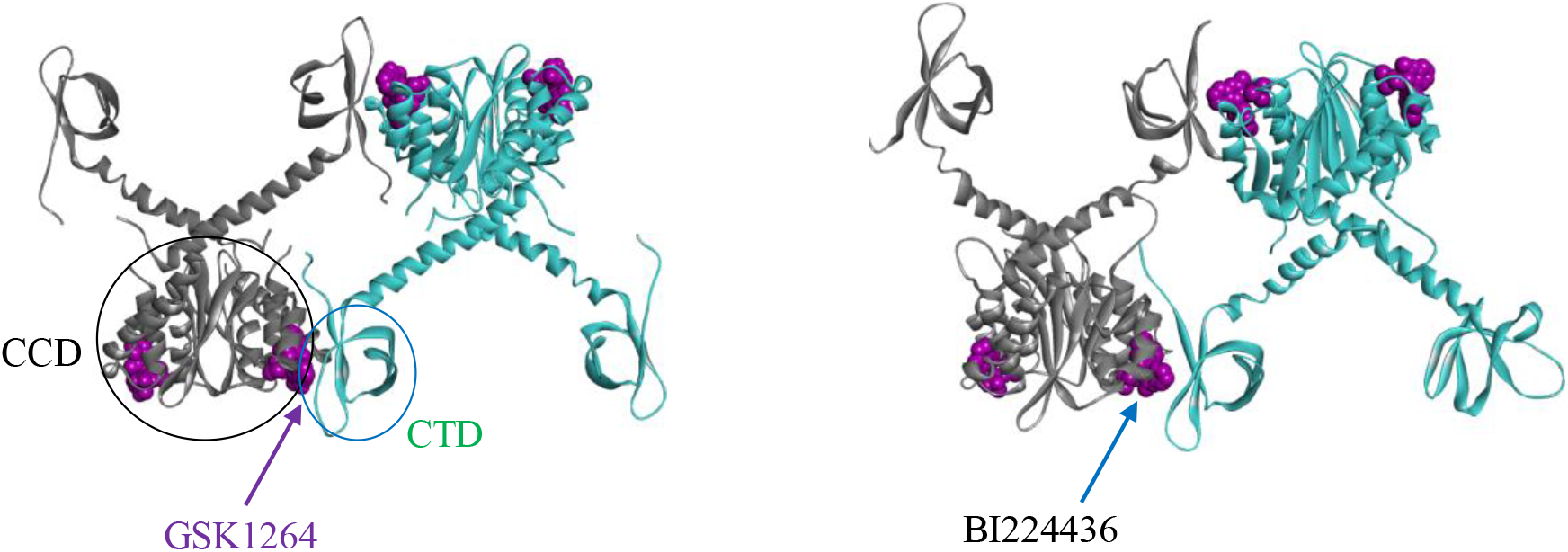
Left: Crystal structure of the IN aberrant tetramer containing the GSK compound GSK1264. Right: A representative MD snapshot taken at 28 ns (starting from two dissociated dimers, each containing the Gilead compound BI224436).

### The probability for observing the native aberrant tetramer within a timescale of hundreds nanoseconds is highly sensitive to initial dimer-dimer separation

These results therefore show that there is a very sharp increase in the mean first passage time (MFPT) to association (Table 1) as the initial separation between the CTD and CCD on the two dimers is increased from about 4.3 Å to 7.8 Å: when the two dimers are separated by a gap of ≤ 4.3 Å, which is less than two water diameters, the association is fast, with MFPT ~ 20 ns for reaching the near native complex (Fig. 3A). Beyond the critical separation gap, the association kinetics is characterized by a much longer association time scale (Fig. 3B) beyond the microsecond range explored in this work.

**Figure 3.**
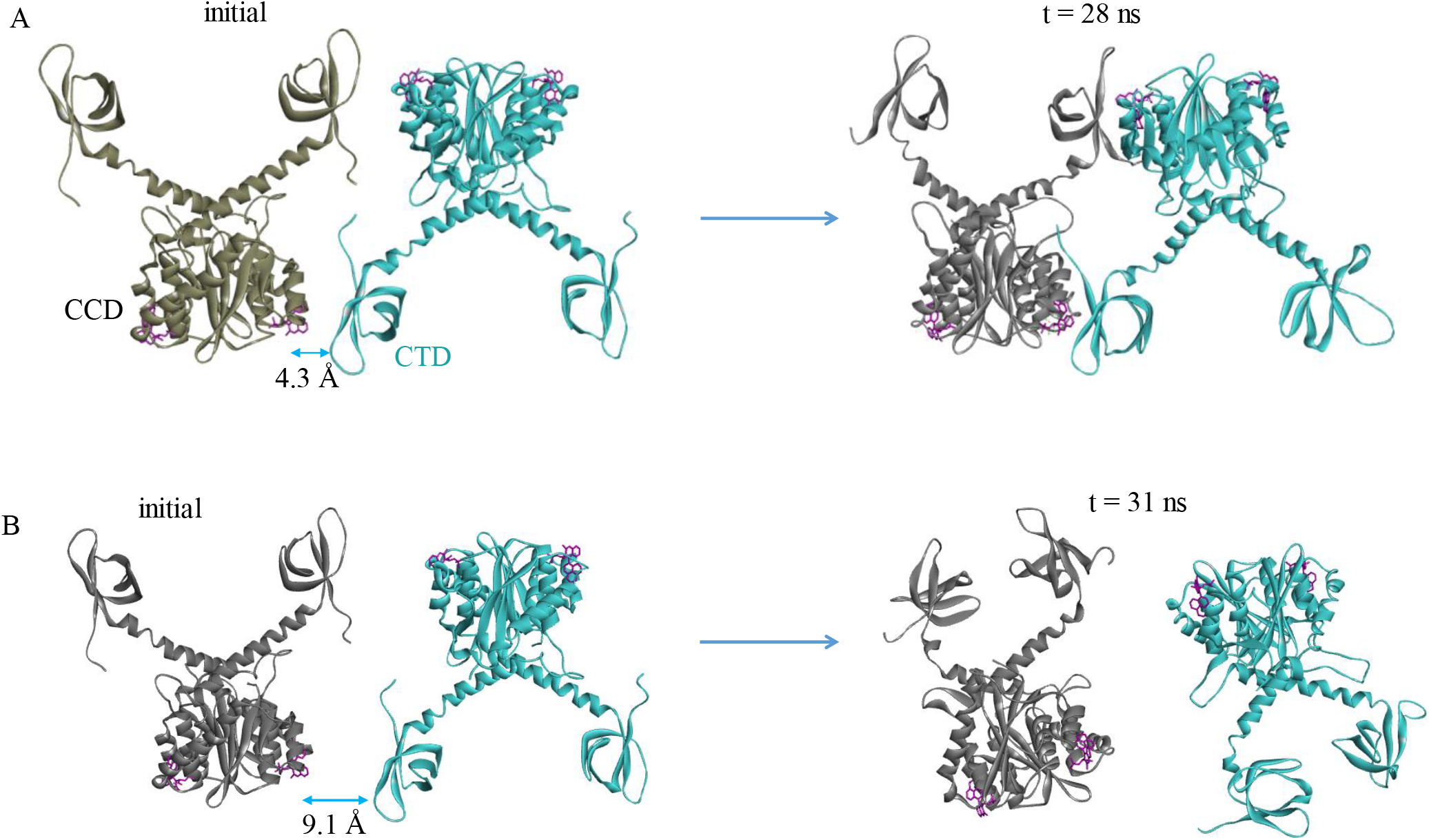
(A) Representative results of simulations Set 1 where the CTD and CCD on the two dimers are initially separated by interfacial gap d ≥ 4.3 Å. (B) A trajectory from simulations Set 2 where the closest distance between CTD and CCD in the starting structure is 9.1 Å.

**Figure 4.**
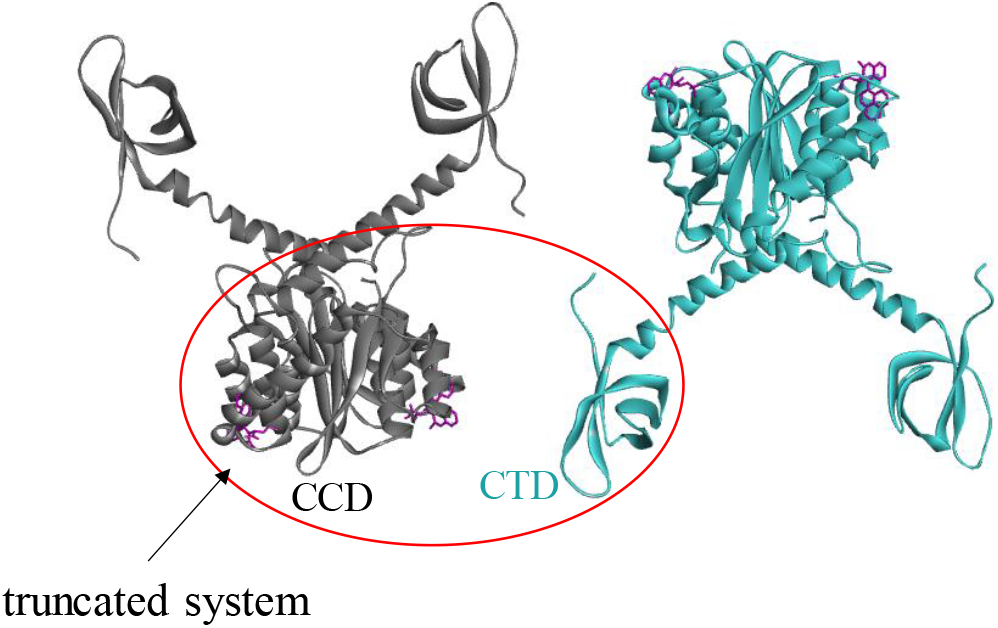
The truncated system (red oval) used to compute the PMF of CTD-CCD association using umbrella sampling.

### The calculated potential of mean force (PMF) reveals that the association free energy landscape is a narrow funnel with the downhill association beginning at a critical separation of about 4.1 Å

To quantitatively characterize the energy landscape, the strength and the molecular mechanism of the drug-induced IN aberrant association, we compute the association free energy profile or potential of mean force (PMF)^13, 28–31^ as a function of the separation distance of the two binding partners. The solvated system of the two dimers has ~180,000 atoms, which makes the PMF calculation computationally very costly. From Fig. 2, it is noted that in the complexed state, two dimers make two symmetrical contact interfaces that each involves one CTD and one CCD dimer, bridged by one drug molecule. Since the two symmetrical contact interfaces are equivalent, a truncated system of one CCD dimer and one CTD (Fig. 5) provides a reasonable model system for investigating the free energy landscape of the association of the full dimer-dimer system. This truncated system (~70,000 atoms) used in the umbrella sampling calculation of the PMF comprises one CCD dimer with a bound ALLINI and one CTD. We show below that the validity of the truncated computational model is supported by the comparison of the onset of the downhill association from the truncated system with that of the full dimerdimer system.

**Figure 5.**
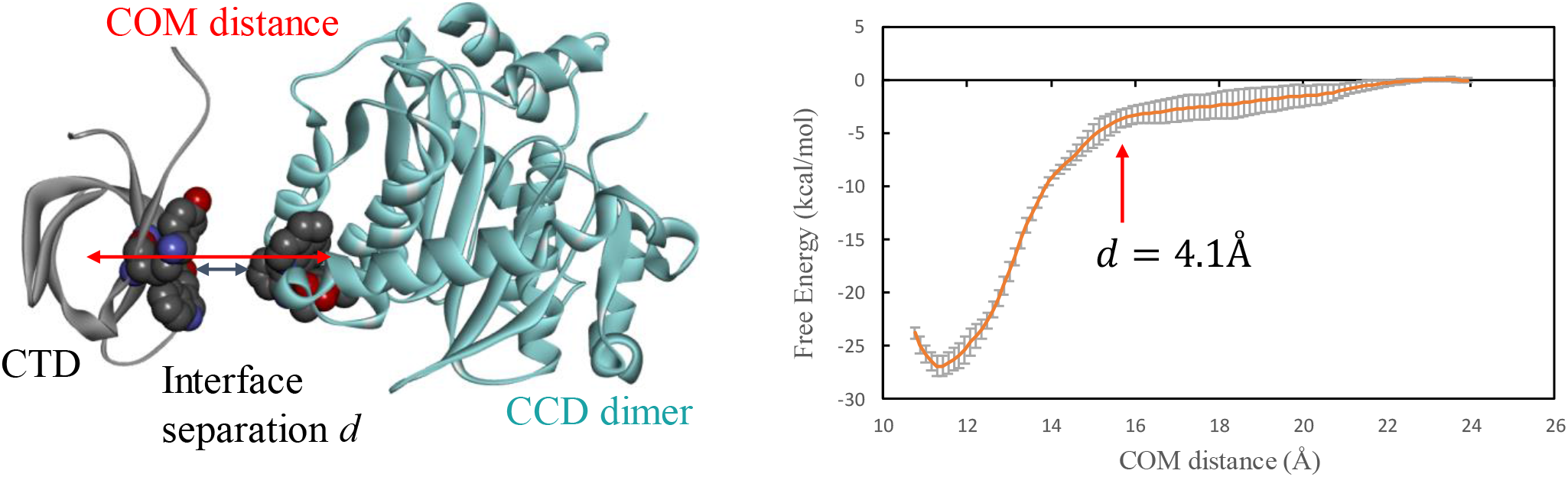
Left: The distance between the center of mass of the CTD and that of the interface residues on the CCD. Also shown is the interface separation distance d between the two binding surfaces. The residues at the centers of the two binding interfaces (Tyr226 and Trp235 on CTD, and BI224436 that binds with CCD) are shown as spheres to illustrate the interface separation d. Right: The PMF of the CTD and BI224436-bound CCD computed using umbrella sampling on the COM distance.

Fig. 5 shows the PMF of the CTD association with an ALLINI-bound CCD dimer, computed from umbrella sampling by applying harmonic bias on the COM distance, i.e. the distance between the center of mass (COM) of the CTD and that of the following interfacial residues on the CCD dimer: 124-128, 131-132, 167-168 and BI224436. The interface separation gap *d* between the two binding surfaces is also shown in the left panel of Fig. 5. Here the gap *d* is calculated by subtracting the COM distance corresponding to the bound complex *d_bound_* = 11.4 Å from the current COM distance. During umbrella sampling the reversible pulling is done along the axis defined by the two COMs defined above, which is perpendicular to the binding interface. This ensures that no intermolecular entanglement or collision between the two binding partners occurs along this pulling pathway. To accelerate the convergence in computing the PMF of protein-protein association, weak angular harmonic restraints on the center of the CTD and its relative orientation with respect to the CCD are applied to the polar angles (θ, ϕ) and three Euler angles (Θ, Φ, Ψ)^23–24^ during umbrella sampling: see Methods.

The PMF is a monotonically decreasing function of the separation. Starting at a critical separation distance of about 4.1 Å the landscape of the protein-protein association becomes a steep downhill funnel. As shown in Fig. 5, the free energy of bringing the two proteins from far apart to *d* = 4.1 Å, i.e. Δ*G*(*d*: ∞ → 4.1 Å) = −3.6 ± 0.86 *kcal/mol*, whereas going from *d* = 4.1 Å to *d* = 0 Å (corresponds to the bound complex), the free energy drop is much larger, Δ*G*(*d*:4.1 Å→ 0 Å) = −23.3 ± 0.43 *kcal/mol*. This result coincides with the observations from the unrestrained molecular dynamics simulations of the full dimer-dimer system, where the spontaneous association is found to occur at an initial separation distance of *d* = 4.3 Å (Table 1 and Fig. 3). The result therefore confirms that the effective attractive interaction involved in the multimerization is very short-ranged, i.e. it becomes attractive at a distance that can accommodate less than two layers of water molecules. Beyond this critical short separation, the association free energy landscape is almost flat and the corresponding association kinetics is largely diffusive. The short-ranged nature of the effective attraction is likely to have contributed to the slow kinetics for multimerization; for example, as shown by the study from Kvaratskhelia group the timescale for the growth of the ALLINI-induced aggregates is on the order of minutes^7^.

### Justification for using PMF alone as a measure of the strength of association

While the strength of molecular association is normally measured by the absolute binding free energy of the two binding partners, for reasons given below, we only need to consider the PMF as a measure for the strength of the protein-protein association. The absolute binding free energy can be written as^23–24^

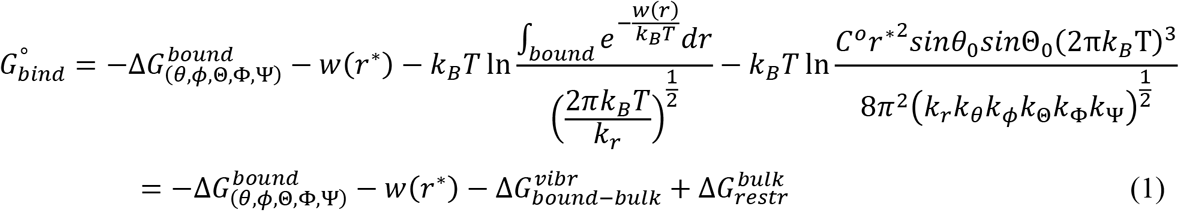

where 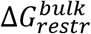 and 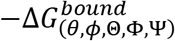 are the free energy of switching on the polar and orientational restraints (*U_θ_, U_ϕ_*, *U*_Θ_, *U*_Φ_, *U*_Ψ_) for the ligand (in this case, CTD) in the bulk solution and that from and switching them off in the bound state, respectively. *w*(*r*\*) is the reversible work (PMF) of pulling the ligand from the bound state to the bulk location *r*\*, computed using umbrella sampling in the presence of the angular restraints (*U_θ_, U_ϕ_*, *U*_Θ_, *U*_Φ_, *U*_Ψ_). 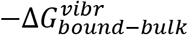 accounts for the ligand positional fluctuation in the bound state versus when it is harmonically restrained at a bulk location with the force constant *k_r_*.^23^ *k_θ_, k_ϕ_*, *k*_Θ_, *k*_Φ_, and *k*_Ψ_ are the force constants for the two polar angles (θ, ϕ) and three Euler angles (Θ, Φ, Ψ) respectively, for the ligand in the receptor reference frame^23–24^. Table 2 lists the different components of the absolute binding free energy for the CTD association with an ALLINI-bound CCD dimer. The calculated 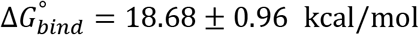 has the same order of magnitude as the experimental affinity for tight protein-protein complexes such as the barnase-barstar (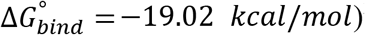^32^.

**Table 2.**
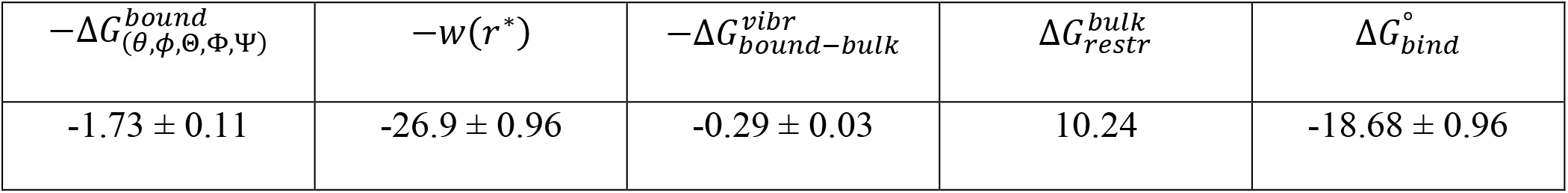
Components of the absolute binding free energy for the CTD association with an ALLINI-bound CCD dimer. (kcal/mol)

| $-\Delta G_{(\theta,\phi,\Theta,\Phi,\Psi)}^{bound}$ | $-w(r^*)$ | $-\Delta G_{bound-bulk}^{vibr}$ | $\Delta G_{restr}^{bulk}$ | $\Delta G_{bind}^\circ$ |
| --- | --- | --- | --- | --- |
| $-1.73 \pm 0.11$ | $-26.9 \pm 0.96$ | $-0.29 \pm 0.03$ | 10.24 | $-18.68 \pm 0.96$ |

As can be seen from Table 2, among the different contributions to the absolute binding free energy 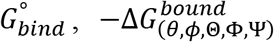 and 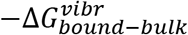 are small compared with the PMF term −*w*(*r*\*). In addition, 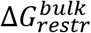 is a constant term for the various combinations of protein-protein association calculations considered in this work. Therefore, in this work we only need to consider the PMF contribution −*w*(*r*\*) as a measure of the absolute association free energy 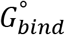.

### Entropy-enthalpy decomposition of the PMF reveals a two-step association process with distinct thermodynamic signatures

To elucidate the thermodynamic driving forces for the association, we decomposed the free energy of association into enthalpic and entropic contributions. Here the free energy of association is taken to be the value at the contact minimum in the PMF near *d* = 0 Å that corresponds to the bound complex relative to the separated state. The entropy of association is estimated from the temperature derivative of the free energy, i.e. 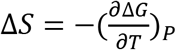, where the Δ*G* is obtained from the PMF at three temperatures 275 K, 300 K and 320 K. As seen from Fig. 6A, the temperature dependence of the free energy of association is approximately linear.

**Figure 6.**
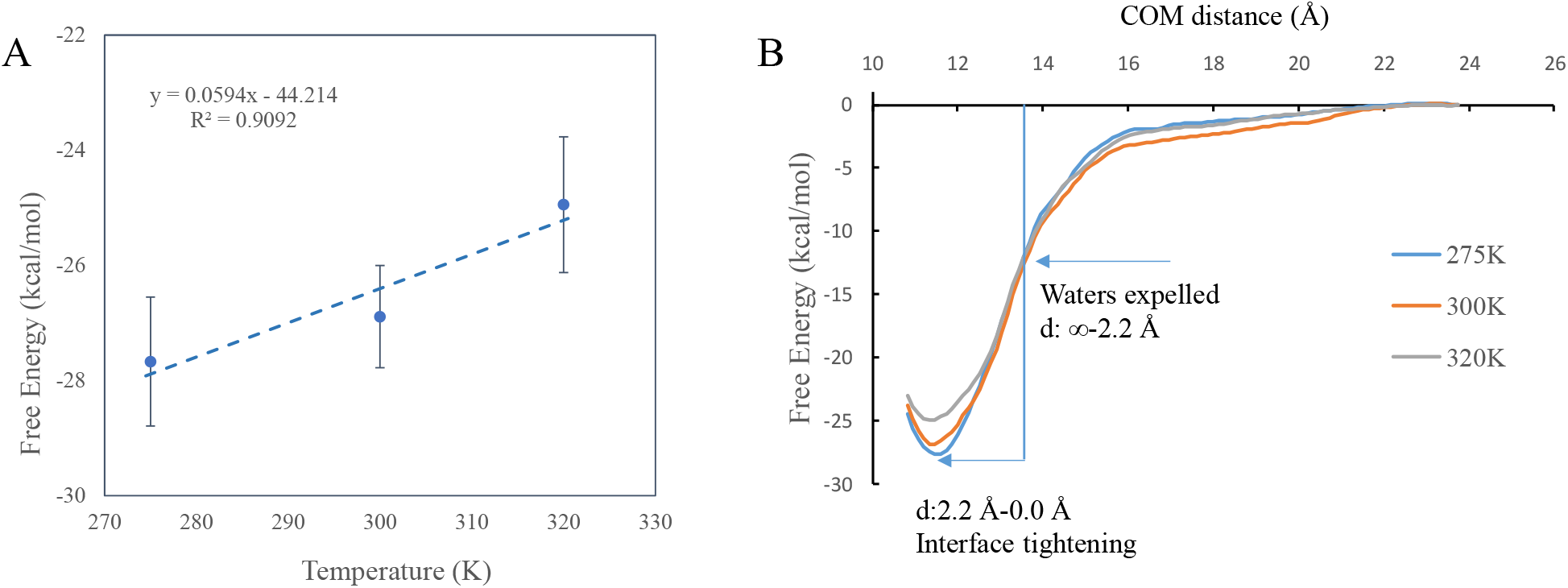
A. Temperature dependence of the free energy of association. B. The range of the PMF can be divided into two parts: the water expulsion phase and the interface tightening phase.

Table 3 gives the thermodynamic quantities of the protein-protein association at different separations between the two binding partners. The column “d: ∞ to 0 Å” shows the thermodynamic quantities for bringing the CTD and the ALLINI-bound CCD from infinitely apart to the contact minimum. This column therefore gives the total association free energy, entropy and enthalpy. As seen from Table 3, overall, the association Δ*G*(∞ → 0 Å) = −26.9 ± 0.96 *kcal/mol* is driven by a large favorable enthalpic contribution Δ*H*(∞ → 0 Å) =−44.7 ± 5.54 *kcal/mol*, and opposed by an unfavorable entropy change −*T*Δ*S*(∞ → 0 Å) = 17.8 ± 5.46 *kcal/mol*. Note that the estimation of the enthalpy and entropy components has much larger statistical uncertainty than that in the free energy, which is expected since the entropy and enthalpy are related to the derivative of free energy^33–34^.

**Table 3.** Thermodynamic properties of the CTD-CCD association at different separations (300 K)

| Thermodynamic quantity | d: ∞ to 0 Å | d: ∞ to 2.2 Å | d: 2.2 Å to 0 Å |
| --- | --- | --- | --- |
| $-T\Delta S$ | $17.8 \pm 5.46$ | $-1.29 \pm 6.08$ | $19.11 \pm 6.03$ |
| $\Delta H$ | $-44.7 \pm 5.54$ | $-10.26 \pm 6.10$ | $-34.45 \pm 6.09$ |
| $\Delta G$ | $-26.9 \pm 0.96$ | $-11.55 \pm 0.48$ | $-15.34 \pm 0.85$ |

Given the fact that protein-protein association is accompanied by releasing water molecules to the bulk solution from the intervening space between the two proteins, the large unfavorable entropy of association is puzzling. To examine the physical origins of the entropic and enthalpic changes, we divide the entire range of the PMF into two segments and compute the corresponding thermodynamic quantities: see Fig. 6B. The first phase consists in bringing the dissociated proteins to a separation of *d* = 2.2 Å where all the water molecules from the intervening space are expelled to the bulk. As shown in Table 3, during this phase the corresponding entropy change is weakly favorable, with −*T*Δ*S*(∞ → 2.2 Å) = −1.29 ± 6.08 *kcal/mol*. This small favorable entropy results from releasing the interfacial water molecules to the bulk, and its small magnitude is attributable to fact that the increase in solvent entropy is largely offset by the loss of side chain entropies as the two binding surfaces approach each other to establish intermolecular contact during this phase. For example, during this phase of the association the Lys266 side chain starts to form a salt bridge with the carboxylate group of the BI224436 at *d* ≈ 3.8 Å. Such intermolecular interactions are also reflected by the favorable Δ*H*(∞ → 2.2 Å) − 10.26 ± 6.10 *kcal/mol*, see Table 3.

The second phase in the association process is from *d* = 2.2 Å until the contact minimum at *d* = 0 Å. This phase is characterized by a small change in the separation distance of just 2.2 Å while the two binding surfaces adapt to each other to optimize their interaction energy, which results in a large reduction in enthalpy Δ*H*(2.2 Å → 0 Å) = −34.45 ± 6.09 *kcal/mol*, which is gained at the expense of a large entropy loss of – *T*Δ*S*(2.2 Å → 0 Å) = 19.11 ± 6.03 *kcal/mol*. Therefore, the large entropy penalty originates from the loss of conformational entropy of the binding surface residues, since all the intervening waters have already been expelled to the bulk in the previous phase d: ∞ → 2.2 Å. Thus, these results indicate that the large unfavorable entropy change for the overall association is caused by configurational entropy loss partially offset by entropic gains due to the expulsion of interfacial water to the bulk.

### The calculated PMF captures the crucial role of ALLINI in promoting IN multimerization

It is known that ALLINI is required for the aberrant IN multimerization^6, 35^. While the low-resolution crystal structure of the IN aberrant tetramer containing an ALLINI provides the structural basis for the multimer^11^, it does not by itself contain information on the strengths of the association with or without ALLINI. Computationally, we have previously used a FEP thermodynamic cycle to compute the relative association free energy to rationalize how different ALLINIs exhibit different potencies in stabilizing the multimer^5^. Here, we use PMF calculation to directly quantify the role of ALLINI in inducing the IN aggregation. Fig. 7 shows the computed PMF of CCD-CTD association with and without the ALLINI BI224436. The PMF of the association without the drug shows a shallow attractive basin that resembles that of a nonspecific protein-protein association. In contrast, the PMF computed with the bound BI224436 features a deep minimum characteristic of specific tight binding. The free energies of association estimated from the PMF in Fig. 7 with and without the drug are −26.9 ± 0.96 *kcal/mol* and −12.4 ±0.7 *kcal/mol*, respectively. The large difference in the association free energy can be understood from the structures of the complexes at their free energy minima in the PMF: see Fig. 8. The CTD-CCD binding interface in the complex containing the drug shows a significantly greater intermolecular atomic packing than the one without the drug. In addition, in the complex without the drug the CTD-CCD interfacial space is filled with several water molecules. These water molecules, which are trapped in a narrow space between nonpolar interfacial residues (e.g. W235, Y226, L102, A128 and W132), form on average significantly fewer number of hydrogen bonds than the number of hydrogen bonds (≈ 3.5) per molecule formed in bulk water. Therefore, these interfacial water molecules in the complex without the drug likely weaken the association free energy. These simulation results provide a physical explanation for the crucial role of ALLINI binding in promoting the IN aggregation.

**Figure 7.**
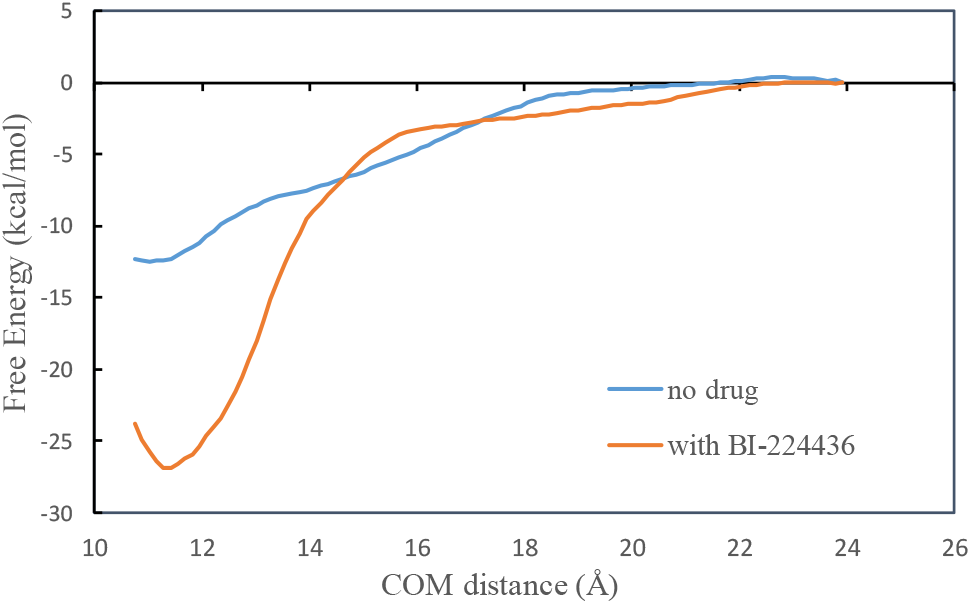
The computed PMF of CCD-CTD association with and without the drug BI224436.

**Figure 8.**
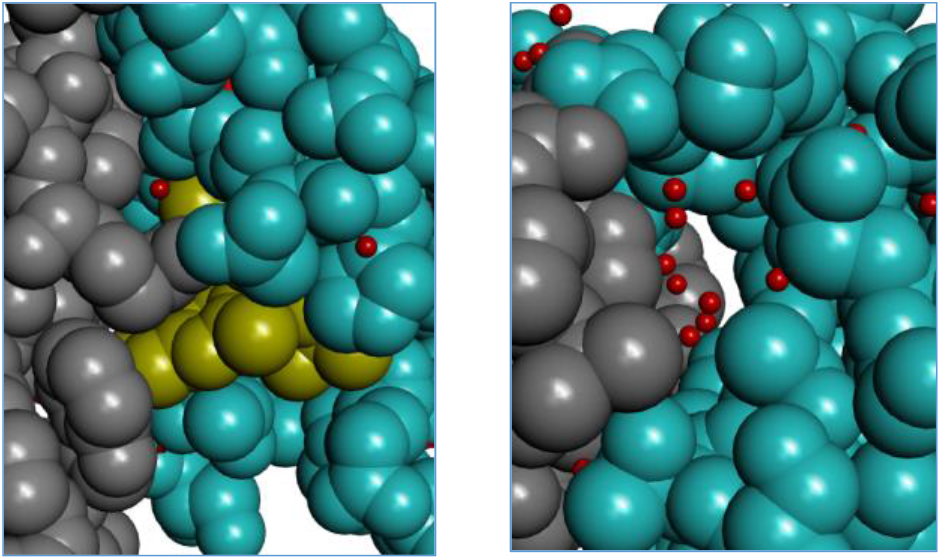
Representative MD snapshots of the complex of CTD (gray) and CCD (cyan) in the presence (left) and absence (right) of BI224436 (yellow). The snapshots are taken from the lowest free energy bins in the PMF (Fig. 7). Water molecules are shown as small red dots.

### The calculated PMF also explains why certain amino acid substitutions exhibited remarkable resistance to the drug-induced IN multimerization

Experimentally a number of engineered amino acid substitutions on the CTD, such as Y226D^11^, W235A^11^ and K266A^7^, are found to abrogate the ALLINI-induced aberrant IN multimerization. While the low-resolution crystal structure of the IN aberrant tetramer shows that these residues are at the protein-protein interface, the experimental structure itself does not give information on the strength of the multimerization from the mutant proteins. Here we run PMF calculations on these mutant proteins to directly probe their effects on the association free energy. Fig. 9 shows the calculated PMF for the three mutant proteins. The calculated association free energies for the three mutant proteins, Y226D, W235A and K266A are −17.5 ± 1.0 kcal/mol, −23.4 ± 2.4 kcal/mol and −20.2 ± 2.7 kcal/mol, respectively. Thus, all three mutants bind significantly more weakly to the CCD than does the wild type CTD (Δ*G_assoc_* = −26.9 ± 0.96 *kcal/mol*), in good agreement with the results of experimental multimerization assays^7, 11^. These results provide further support for the validity of the free energy model adopted in this work.

**Figure 9.**
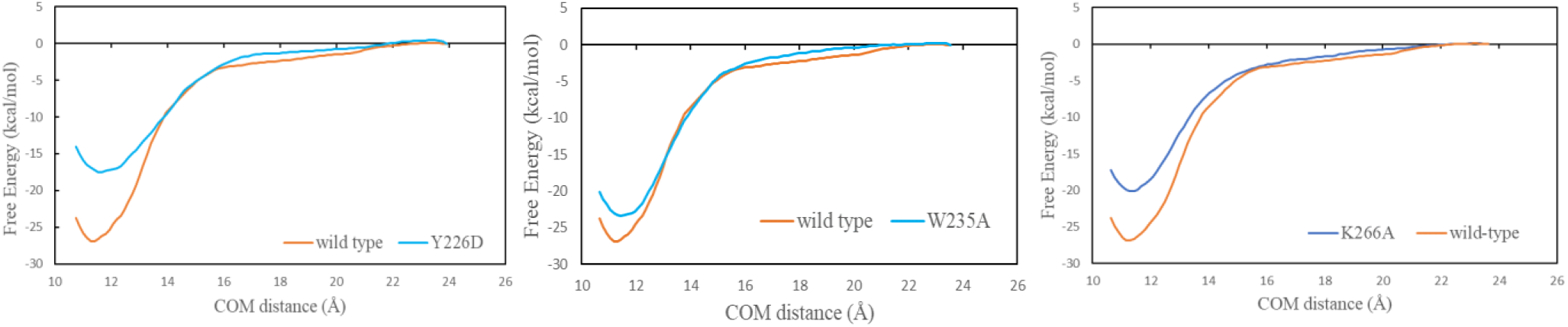
Free energy profile of the CTD-CCD binding in which the CTD contains the following single mutations: Y226D (left), W235A (center) and K266A (right).

## Discussion

We have applied direct MD simulations and free energy calculations to explore the energy landscape of the protein-protein interaction responsible for the drug-induced aberrant multimerization of HIV-1 integrase. The PMF results captured the critical role of the ALLINI in promoting the CTD-CCD intersubunit interactions that are the cornerstone of the IN aggregation (Fig. 7 and Fig. 8). By showing that all three key substitutions Y226D, W235A and K266A results in significantly less attractive association than the wild type protein (Fig. 9), the calculations also provide physical explanations for why these amino acid substitutions block the drug-induced IN multimerization. These results demonstrate the power of free energy modeling in elucidating the effects of mutations in protein-protein interactions. However, potential of mean force calculations in explicit solvent as those performed here remain computationally very demanding. In this regard we note a recent study by Siebenmorgen & Zacharias who have used a PMF-based binding free energy method in an implicit solvent to more accurately re-rank the docking poses of 20 different protein-protein complexes^15^.

In addition to providing a physical basis for the IN multimerization, the work sheds light on the thermodynamic foundation of protein-protein association. We find that the protein-protein association between the CTD and CCD-ALLINI is driven by enthalpy change, and opposed by entropy: see Table 3 and Fig. 6. (Note that in the following we focus on the solvent and solute conformational entropy changes only, which are the values given in Table 3; the external entropy change that includes translational and rotational entropies will not be discussed.) To examine the origin of the favorable entropy we compare the free energy profile for the CTD and CCD-ALLINI association with the direct interaction energies that make up the total enthalpy of dimerization: the protein-protein, protein-solvent, and solvent-solvent interaction (Fig. 10). Although all the interaction energies are much noisier compared with the well-behaved free energy profile, it can be seen that as the two proteins approach each other, the association is favored by the protein-protein interaction energy. Conversely, the association is strongly opposed by the loss of protein-solvent interactions that are replaced by protein-protein interactions. Solvent-solvent interactions favor dimerization because water molecules expelled from the inter-protein space form more hydrogen bonds in the bulk. Note that the solvent-solvent interaction and the proteinprotein direct interaction have similar orders of magnitudes.

**Figure 10.**
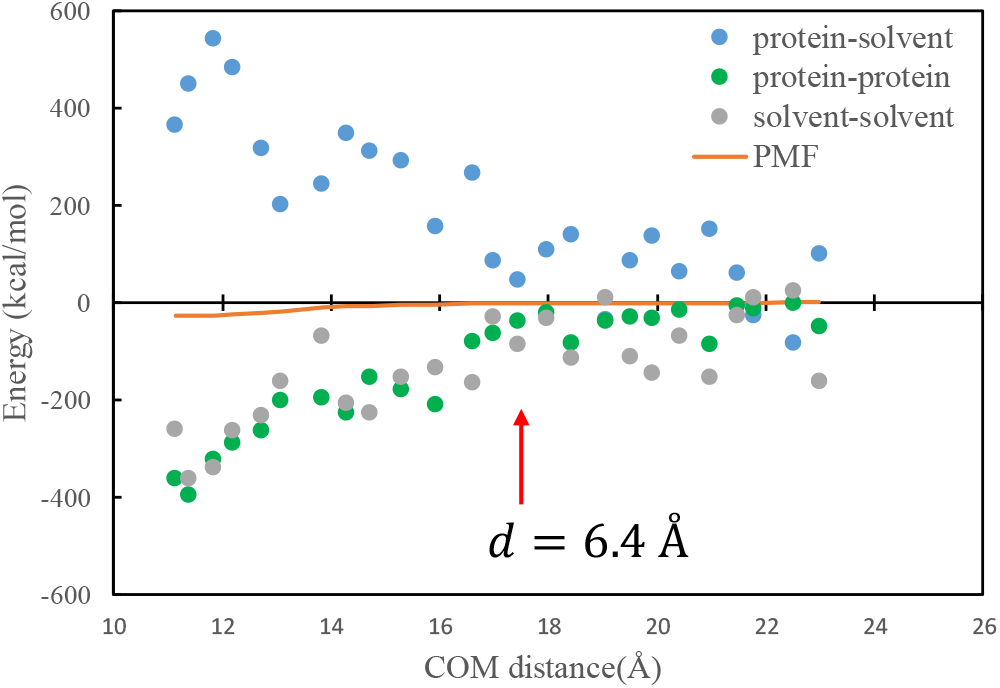
The PMF or free energy (orange) plotted together with the various interaction energies: protein-protein (green), solute-solvent (blue) and solvent-solvent (gray).

We now discuss these results in the context of hydrophobic and hydrophilic protein-protein association, which can have different thermodynamic signatures. It is has been known that the thermodynamic driving force for hydrophobic association is length scale dependent^36–37^. Aggregation of small hydrophobic solutes is driven by favorable entropy change and opposed by unfavorable enthalpy change. This is because the water molecules near small hydrophobic solute can rearrange themselves to maintain the hydrogen bond network and, therefore, releasing such surface waters to the bulk upon solute association is more or less enthalpy neutral. In this case, the net enthalpy change of association results mainly from the loss of favorable solute-water interactions, which can outweigh the favorable solute-solute interaction. For example, Cui et al.^38^ studied the thermodynamics of the rigid body binding between a hydrophobic α-helix with ubiquitin. The size and shape of the buried surface area in this complex suggest that it belongs to the small solute regime. Hence the binding is driven by entropy and opposed by enthalpy, due to the loss of protein-water interactions not compensated by the gain of protein-peptide interactions and solvent-solvent interactions from the desolvation of the solutes^38^.

For large hydrophobic solutes (e.g. interface dimension *R* > 10 Å) association can be favored or disfavored by enthalpy, depending on the strength of the solute-water interaction. This is because unlike the water molecules around a small solute, water molecules near a large hydrophobic surface cannot orient themselves around the surface to maintain the hydrogen bond network. As a result, the loss of large hydrophobic surface patches upon binding releases energetically frustrated waters molecules into the bulk resulting in a reduction of the enthalpy. For example, using highly hydrophobic plates with weak solute-water interaction parameters, Zangi and Berne^39^ found that both entropy and enthalpy favor hydrophobic collapse. However, for graphene plates that have stronger plate-water interaction parameters, Choudhury and Pettitt^40^ found that the collapse of the plates in water is driven only by entropy and opposed by enthalpy. This is because the loss of plates-water interaction upon collapse outweighs the enthalpy gain from the release of water molecules in between the plates and from the increased direct plate-plate attraction.

There have been few computational studies on characterizing the thermodynamic driving forces for hydrophilic protein-protein association^41–42^. Helms and coworkers have studied several hydrophilic protein-protein complexes including the barnase-barstar complex^41^, focusing on the calculation of the binding free energy and translation/rotation entropy changes. For the barnase-barstar complex, using the experimental data^43^ one can estimate that the protein-protein binding is largely driven by favorable enthalpy with a relatively small favorable entropy contribution. Ben-Naim proposed a model suggesting that in hydrophilic protein-protein association, water molecules can “stitch” the two proteins by forming hydrogen-bond bridges between two hydrophilic groups on the surfaces of two binding partners, resulting in favorable enthalpy at the expense of entropy^44^.

In the present study, as shown in Fig. 11 the protein-protein interfaces in the complex contain a mixture of mainly hydrophobic groups and also several hydrophilic residues, with the nonpolar patch roughly surrounded by polar residues. Since both the solvent and solute degrees of freedom are unconstrained, our calculations reveal a more realistic thermodynamic picture of protein-protein association than did previous studies which either used idealized geometry (plates and ellipsoids) or have fixed intramolecular degrees of freedom of the protein. Importantly, we found that when the protein conformational degrees of freedom are accounted for, the entropy of association becomes unfavorable (Table 3). Consistent with previous work, we also found that during the water expulsion phase, the entropy change is weakly favorable, as the large increase in the solvent entropy due to inter-protein water release is largely cancelled by the reduction in the solute entropy. As the two proteins approach the contact minimum, the entropy drops further as the conformational motions are more limited. The configuration entropy loss at the final stage of the complexation of ~ 20 kcal/mol is in the same order as those estimated by Chang et al. for the binding of AKAP protein HT31 with the D/D domain of RII α-regulatory subunit of protein kinase A^45^.

**Figure 11.**
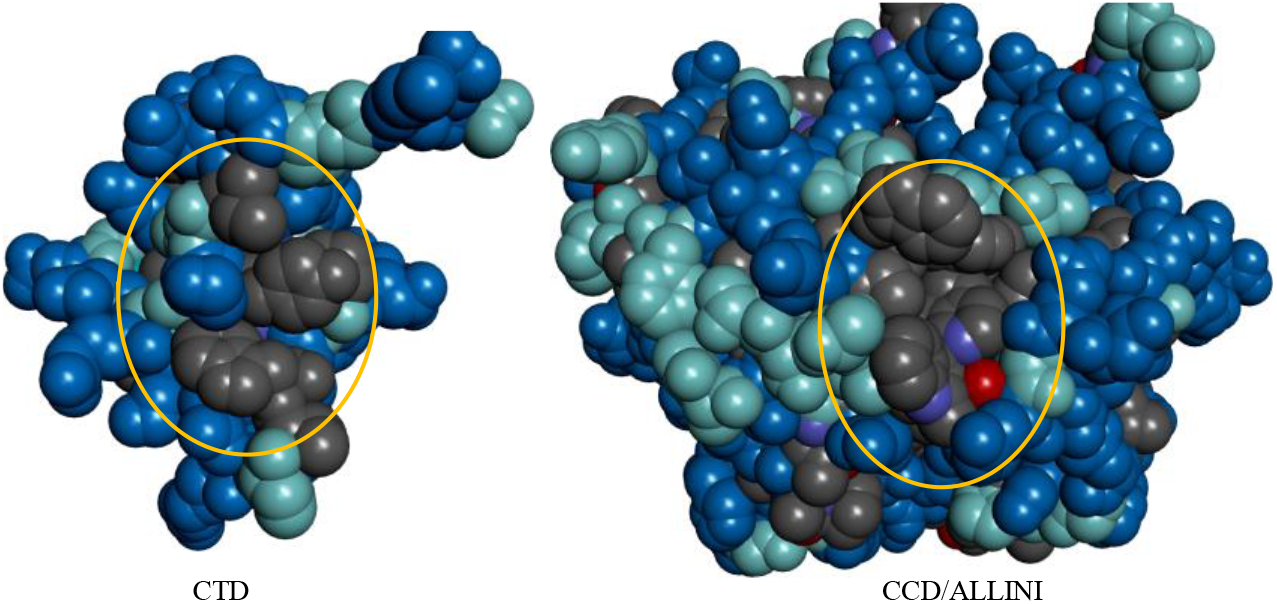
The VDW surfaces of the binding interfaces in the two proteins. Gray: hydrophobic. Blue: hydrophobic. Cyan: intermediate between hydrophilic and hydrophobic. The central contact residues involved in the binding are circled.

In the current study, we find evidence that is consistent with the bridging water effect proposed by Ben-Naim: see Fig. 12. The thermodynamic properties for the association of large hydrophobic and hydrophilic solutes surveyed here are summarized in Table 4. It should be noted that while the signs of the entropy and enthalpy may be generalized for idealized systems, for realistic proteins the situation is more complex and likely to be system dependent because of the presence of many variables such as the interface composition, curvature, flexibility, and solvent conditions (e.g. ionic concentration and pH).

**Figure 12.**
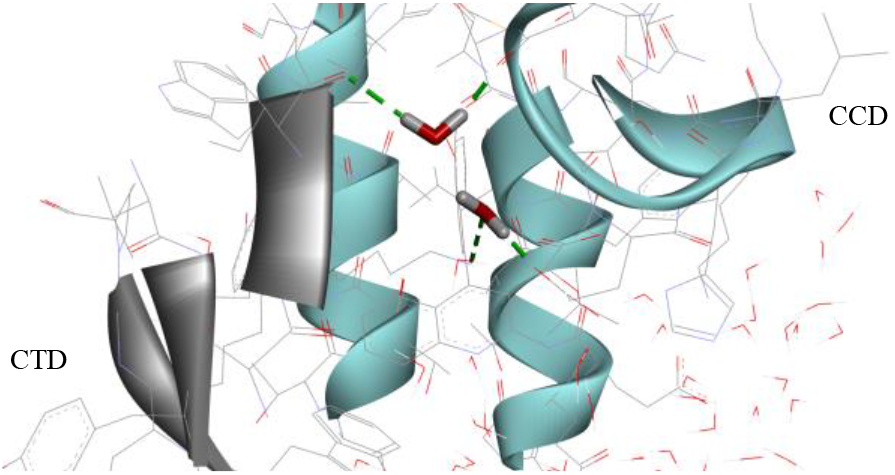
A MD snapshot showing two bridging waters forming hydrogen bonds with both proteins.

**Figure 13.**
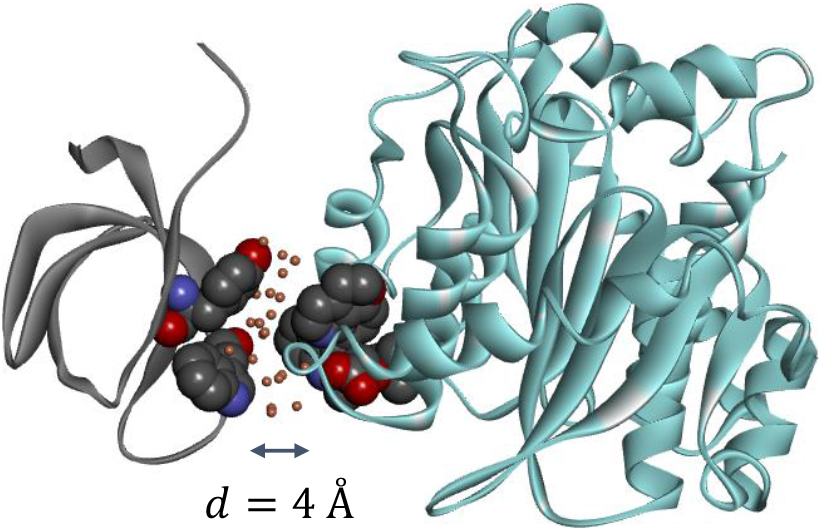
The orange dots represent interfacial water sites whose occupancy is at least 1.5 times the bulk water occupancy.

**Table 4.**
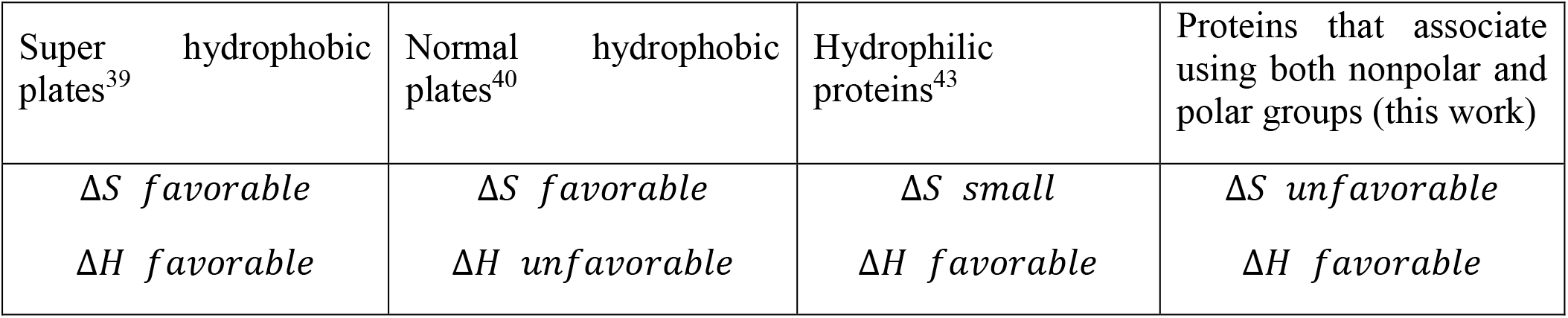
Summary of the thermodynamic driving forces in the association of large solutes surveyed here

In this work, both the direct simulations and PMF calculations show that the effective range of the attraction between the two binding proteins CTD and ALLINI-bound CCD is very short, at interface distance *d* ≈ 4 Å or less than two layers of water molecules, even though the two binding partners CTD and CCD carry 6+e and 3-e net charges, respectively. Apparently, the short-ranged effective interaction is the combined result of several factors, including the spatial distribution of the charges and the electrostatic screening from the solvent. To gain more insights into the origin of this short-ranged effective interaction we analyze the calculated protein-protein PMF in terms of the interaction energy components in Fig. 10. Although all the interaction energies are much noisier compared with the well-behaved free energy profile, some qualitative trends can be gleaned from such an analysis. First, the short-range proteinprotein PMF is the result of near complete cancellation of several much larger forces (note that entropy contributions not included in Fig. 10). Second, the interaction energies have somewhat longer range compared with the free energy. For example, the protein-solvent interaction energy starts to increase at about *d* = 6.4 Å, while the free energy starts its descent at *d* = 4.1 Å.

Next, we compare the effective range of the protein-protein binding observed in this work with those reported in the literature^13, 41–42, 46^. Helms and coworkers have computed the PMF for several hydrophilic protein-protein complexes including the barnase-barstar complex^41^ where they report a large effective range of the binding funnel of ≈ 14 Å. The PMF of the barnase-barstar complex, which is characterized by highly polar/charged interface in explicit solvent, has also been reported by other groups. For example, Hoefling and Gottschalk^46^ reported the width of the binding funnel to be ~15 Å, similar to the result by Helms and coworkers^41^. On the other hand, in a study by Gumbart et al^13^ in which the protein conformational degrees of freedom are constrained, the effective range of attraction in barnase-barstar is ~6 Å. Overall, the effective range of the association funnels for hydrophilic protein-protein binding is considerably larger than the ~4 Å for the HIV-1 integrase aggregation studied here. The difference in the width of the binding funnel could be attributed to the more hydrophobic nature of the system studied in the present work, as the CTD and drug-bound CCD interface involve several nonpolar residues such as A128, T226, W235 and I268, in addition to the charged K266.

Lastly, some studies of hydrophobic protein-protein association^37, 47–49^ have focused on determining the effects of dewetting or cavitation^50^, i.e. the cooperative liquid-to-vapor phase transition in the interdomain region which was first observed in idealized plates when the inter-plate region is still large enough to hold more than two layers of water.^37^ While a dewetting transition may occur below a critical separation between idealized nanoscale hydrophobic plates or ellipsoids^51–52^, it has rarely been observed in realistic hydrophobic protein association^53^. For example, in the hydrophobic collapse of protein domains of 2,3-dihydroxybiphenyl dioxygenase (BphC), dewetting is observed only after turning off the protein-water electrostatic interactions in the simulation^48^. The only system in which the cooperative drying phase transition has been observed is in the melittin tetramer^47^, which was separated to create a nanoscale tube-like channel. Interestingly, Ricci and McCammon^54^ recently observed that during the association of a 15-residue hydrophobic MDM2 peptide to a highly hydrophobic cleft of the p53 protein, the water density in the interfacial space starts to drop from the bulk value when the interfacial separation is less than 7.6 Å, which is believed to be the onset of dewetting transition. However, it appears that cavitation in the inter-protein space did not occur even in this highly nonpolar system. In the present study, we did not observe a dewetting transition in the protein-protein association of the CTD and CCD-ALLINI system. As shown in Fig. 12, within the binding funnel, the intervening region remains well hydrated with water molecules whose occupancy is even higher than that in the bulk water. The small number of hydrophilic groups in the interface region (Fig. 11) which interact strongly with interfacial water molecules is likely cancelling any tendency to dewet^53^.

## Conclusion

Elucidating the energetics and thermodynamic aspects of protein-protein association is important to our understanding of this fundamental molecular recognition process. Computational characterizations of the thermodynamic signatures of protein-protein association for realistic systems are still scarce. Here using explicit solvent MD and umbrella sampling, we have provided a more complete picture of the proteinprotein association responsible for the drug-induced HIV-1 integrase multimerization. The calculated free energy profile has correctly captured the critical role of the inhibitor in the integrase multimerization, and accounted for the effects of various amino acid substitutions on the protein association. Remarkably, the results reveal that the effective range of the protein-protein attractive funnel is less than two layers of water molecules, which is significantly shorter than those reported for the more hydrophilic proteinprotein complexes, despite the fact that the two binding partners studied here carry opposite net charges. Lastly, we find that while the majority of the residues forming the contact interface are hydrophobic, the overall association is favored by enthalpy and opposed by entropy. Our analysis suggests that while the solvent release from the inter-protein region into the bulk solution results in an increase in entropy, this is more than offset by the loss of the solute configurational entropy due to the complexation.

## Author Contributions

N. Deng, R. Levy and E. Gallicchio designed the research. C. Tse, L. Wickstrom and N. Deng performed the research. M. Kvaratskhelia contributed research data. N. Deng, R. Levy, E. Gallicchio and L. Wickstrom analyzed the results. N. Deng, R. Levy and E. Gallicchio wrote the paper.

## Acknowledgments

The calculations were run on the XSEDE allocation resource TG-MCB100145 and a shared computing cluster at Temple University supported by NIH S10 OD020095. N. Deng thanks Prof. Wei Yang at Florida State University and Mr. Eric Chen from Lehman College, CUNY for helpful discussions. This study was supported by NIH grant 5U54AI150472-09 and by a Bridge fund from Pace University to N. Deng.

